# Diverse transcriptomic effects of evolved imipenem-relebactam resistance in *Pseudomonas aeruginosa*

**DOI:** 10.64898/2026.09.12.751187

**Authors:** Adeline Supandy, Ava J. Dorazio, Kevin M. Squires, Ellen G. Kline, Ryan K. Shields, Daria Van Tyne

**Affiliations:** Department of Medicine, Division of Infectious Diseases, University of Pittsburgh, Pittsburgh, PA, USA; Antibiotic Management Program, University of Pittsburgh Medical Center, Pittsburgh, PA, USA; Center for Innovative Antimicrobial Therapy, University of Pittsburgh, Pittsburgh, PA, USA; Center for Evolutionary Biology and Medicine, University of Pittsburgh, Pittsburgh, PA, USA

**Keywords:** Imipenem-relebactam, *pseudomonas aeruginosa*, RNA-sequencing

## Abstract

Imipenem-relebactam (Imi/Rel) is a β-lactam/β-lactamase inhibitor combination used for the treatment of multidrug-resistant *Pseudomonas aeruginosa* infections. We previously reported that treatment-emergent resistance to Imi/Rel is associated with mutations in the AmpC beta­lactamase and/or the MexAB-OprM and MexEF-OprN efflux operons. However, the impact of these mutations on bacterial gene expression has not been explored extensively, particularly among *P. aeruginosa* from patients treated with Imi/Rel. To determine the effect of treatment-emergent Imi/Rel resistance on *P. aeruginosa* global transcription, we performed RNA sequencing on paired *P. aeruginosa* clinical isolates from six patients collected before and after Imi/Rel treatment. Transcriptional responses varied substantially, with no conserved changes in gene expression identified across all six patients. Three isolate pairs showed significant upregulation of previously characterized Imi/Rel resistance-associated genes in the treatment-emergent resistant isolate, while two resistant isolates displayed significant downregulation of *ampC*. Comparisons of the top 10 differentially regulated genes in the resistant isolate from each patient revealed only one gene that was shared between all six patients. Pathway enrichment analysis using Clusters of Orthologous Genes (COG) categories suggested that Imi/Rel exposure impacts transcription of genes involved in translation and metabolism, but these changes are highly variable between patients and isolates. Overall, we find that the transcriptional response of clinical *P. aeruginosa* to Imi/Rel exposure appears to be highly diverse and likely dependent on the genetic background of the infecting *P. aeruginosa* strain.

## Introduction

*Pseudomonas aeruginosa* is a Gram-negative pathogen responsible for an increasing number of multidrug-resistant (MDR) hospital-acquired infections (1, 2). The preferred treatment for MDR *P aeruginosa* infections are β-lactam/β-lactamase inhibitor combinations such as ceftolozane-tazobactam and ceftazidime-avibactam (3). Because resistance to these combinations continues to rise (4), a newer combination containing imipenem and relebactam (Imi/Rel), has seen increased clinical usage (5, 6). Resistance to β-lactam/β-lactamase inhibitor combinations can be mediated through upregulation of multidrug efflux systems like MexAB-OprM, loss of OprD porin function, and mutations affecting the chromosomal AmpC [3-lactamase or antibiotic targets like Penicillin-Binding Protein 3 (encoded by *ftsI*) (2, 5, 7).

Unfortunately, Imi/Rel use has resulted in treatment-emergent resistance in some clinical *P. aeruginosa* isolates. We and others previously found that Imi/Rel resistance was associated with mutations affecting the MexAB-OprM and MexEF-OprN efflux operons, either directly or indirectly through mutations in operon regulators (5, 8–10). Mutations affecting the CreBC two-component regulatory system and AmpC expression modulation have also been implicated (5). Whether these mutations act primarily at the transcriptional level, through structural changes to the encoded proteins, or both, remains unclear. As such, further investigation into transcriptional responses associated with Imi/Rel resistance and the functional pathways affected remain to be explored.

In this report, we characterized the transcriptional consequences of treatment-emergent Imi/Rel resistance using clinical isolates of MDR *P. aeruginosa* from six patients with MDR *P. aeruginosa* infections. We generated RNA-seq datasets for each pair of baseline and post­exposure isolates and compared expression profiles within and between patients.

## Methods

### Isolate Collection, DNA Sequencing and Analysis

*P. aeruginosa* clinical isolates were collected from six patients pre- and post-exposure to Imi/Rel for a total of 12 isolates. Genomic DNA from the PAO1 laboratory strain and the six baseline isolates was extracted from overnight cultures grown in cation-adjusted Mueller-Hinton broth at 37°C using a DNeasy Blood and Tissue Kit (Qiagen) according to the manufacturer’s protocol, with the Proteinase K incubation step extended to 60 minutes. Libraries were prepared (2 × 150 bp, paired-end reads) and whole genome sequencing was performed on the Illumina platform at SeqCenter (Pittsburgh, PA). Genomes were *de novo* assembled using SPAdes v3.15.5 and annotated using Prokka v1.14.5 (11, 12). A gene presence absence file was generated using Roary v3.13.0 and used to construct a midpoint-rooted single-copy core genome phylogenetic tree using RAxML v8.2.12 with the GTRGAMMA flag which was visualized in iTOL (13, 14). Multi-locus sequence types (STs) were determined using the PubMLST database (15).

### RNA Sequencing and Analysis

RNA was extracted from all 12 pre- and post-Imi/Rel exposure isolates and was sequenced as previously described (16). Briefly, 5 mL of re-inoculated cultures at 1:100 dilution from overnight cultures were grown to mid-log phase (OD600 0.3 – 0.4) in cation-adjusted Mueller-Hinton broth at 37°C prior to extraction using the RNeasy Mini Kit (Qiagen) according to the manufacturer’s protocol. Total RNA was treated with DNase and 2×150 bp paired-end libraries were generated and sequenced on an Illumina NovaSeq X Plus at SeqCenter (Pittsburgh, PA). Four biological replicates were collected for each isolate, and only R1 reads were used for analysis.

Sequenced reads were trimmed using TrimGalore v0.6.10 (17) and only reads with Phred scores >33 and sequence lengths >142 bp were retained for analysis. Trimmed reads were mapped to Prokka-annotated baseline or PAO1 reference genomes using STAR v2.7.11b with – outFilterScoreMinOverLread and –outFilterMatchNminOverLread flags set to 0.8 (18). Reads mapping to coding sequences were counted using featureCounts v2.0.6 (19). Analysis of differentially expressed genes (DEGs) was conducted using DESeq2 in RStudio (20). Only genes with >10 read counts were included in the analysis. Significantly differentially expressed genes were defined as genes with adjusted *P*-values <0.05 (Benjamini-Hochberg corrected) and log_2_ fold-change <-1 or >1. For principal component analysis (PCA), data were normalized with the varianceStabilizingTransformation() function. Volcano plots were constructed using the EnhancedVolcano package. For analysis of all patient isolates against PAO1, the design parameter of DESeq2 was set to “∼0 + condition”.

### COG Category Enrichment

The function of each differentially expressed gene was determined by assigning Clusters of Orthologous Genes (COG) categories using EggNOG-mapper (21). The number of annotated genes assigned to at least one COG category for PAO1 and each baseline isolate was: 92.3% (5179/5494) for PAO1, 91.9% (5557/6050) for PT1, 93.0% (5533/5952) for PT2, 96.5% (5543/5744) for PT3, 92.2% (5733/6220) for PT4, 94.3% (5164/5474) for PT5, and 93.9% (5070/5397) for PT6 (**File S1)**. Genes with multiple COGs were counted once in each category for analysis. DEGs without an assigned COG category were excluded from analysis, and significance was determined using Fisher’s exact test with Bonferroni correction for multiple testing.

## Results

The median age of the six patients in this study was 55 years, 83% were male, and 50% had a history of solid organ transplantation (**Table 1**). Each patient developed a recurrent MDR *P. aeruginosa* infection that were treated with Imi/Rel for a median (range) of 20 (10–82) days prior to the development of Imi/Rel resistance. We first evaluated the transcriptomes of all 12 isolates against PAO1 to compare bacterial transcriptomes between patients. Baseline isolates from each patient were genetically distinct (**Fig S1**), and the transcriptional profiles of isolates from different patients were largely distinct from each other, except for PT1/PT2 and PT5/PT6, whose transcriptomes clustered together on a principal component analysis (PCA) plot of transcriptomes from all isolates (**Fig 1A**). Furthermore, the profiles of post-exposure Imi/Rel-resistant isolates from each patient closely clustered with their baseline susceptible counterparts, suggesting less within-patient variation in gene expression compared with between-patient variation (**Fig 1A**). We next compared the transcriptomes of all baseline isolates against all resistant isolates using the PAO1 genome as a reference, and identified several significantly differentially expressed genes (DEGs) (**Fig 1B**). Significantly downregulated genes included a hypothetical protein (PAO1_02342), the CTP synthase *pyrG*, and the phenazine biosynthesis gene *phzB1*, while significantly upregulated genes included the flavohemoprotein gene *hmp*, transcriptional regulatory gene *qseB*, and the polyamine aminopropyltransferase gene *speE* (**File S1**). While these data suggest that treatment-emergent Imi/Rel-resistant *P. aeruginosa* isolates might have a conserved transcriptional response, none of the DEGs include previously described Imi/Rel resistance genes (5).

**Table 1.** Patient Demographics and *P. aeruginosa* Imi/Rel MICs.

| Patient | Age | Sex | Underlying disease | MDR *P. aeruginosa* infection | Duration of Imi/Rel treatment prior to resistance (Days) | Isolate type | Imi/Rel MIC (mg/L)* |
| --- | --- | --- | --- | --- | --- | --- | --- |
| PT1 | 56 | M | HIV | Recurrent pneumonia | 20 | Baseline | 1 |
|  |  |  |  |  |  | Resistant | 8 |
| PT2 | 55 | M | Lung transplant | Recurrent pneumonia | 10 | Baseline | 0.5 |
|  |  |  |  |  |  | Resistant | 8 |
| PT3 | 80 | M | Esophageal strictures | Pneumonia and abdominal wall abscess | 12 | Baseline | 1 |
|  |  |  |  |  |  | Resistant | 32 |
| PT4 | 55 | M | Liver transplant | Recurrent pneumonia | 23 | Baseline | 1 |
|  |  |  |  |  |  | Resistant | 8 |
| PT5 | 54 | F | Non-ischemic cardiomyopathy s/p LVAD<sup>#</sup> | Recurrent bacteremia and driveline infections | 82 | Baseline | 0.5 |
|  |  |  |  |  |  | Resistant | 8 |
| PT6 | 72 | M | Heart transplant | Recurrent pneumonia complicated by empyema | 20 | Baseline | 4 |
|  |  |  |  |  |  | Resistant | 32 |

\*Susceptibility testing was performed by broth microdilution in triplicate using a fixed
concentration of relebactam (4mg/L).
<sup>#</sup>LVAD: Left-Ventricular Assist Device

**Figure 1.**
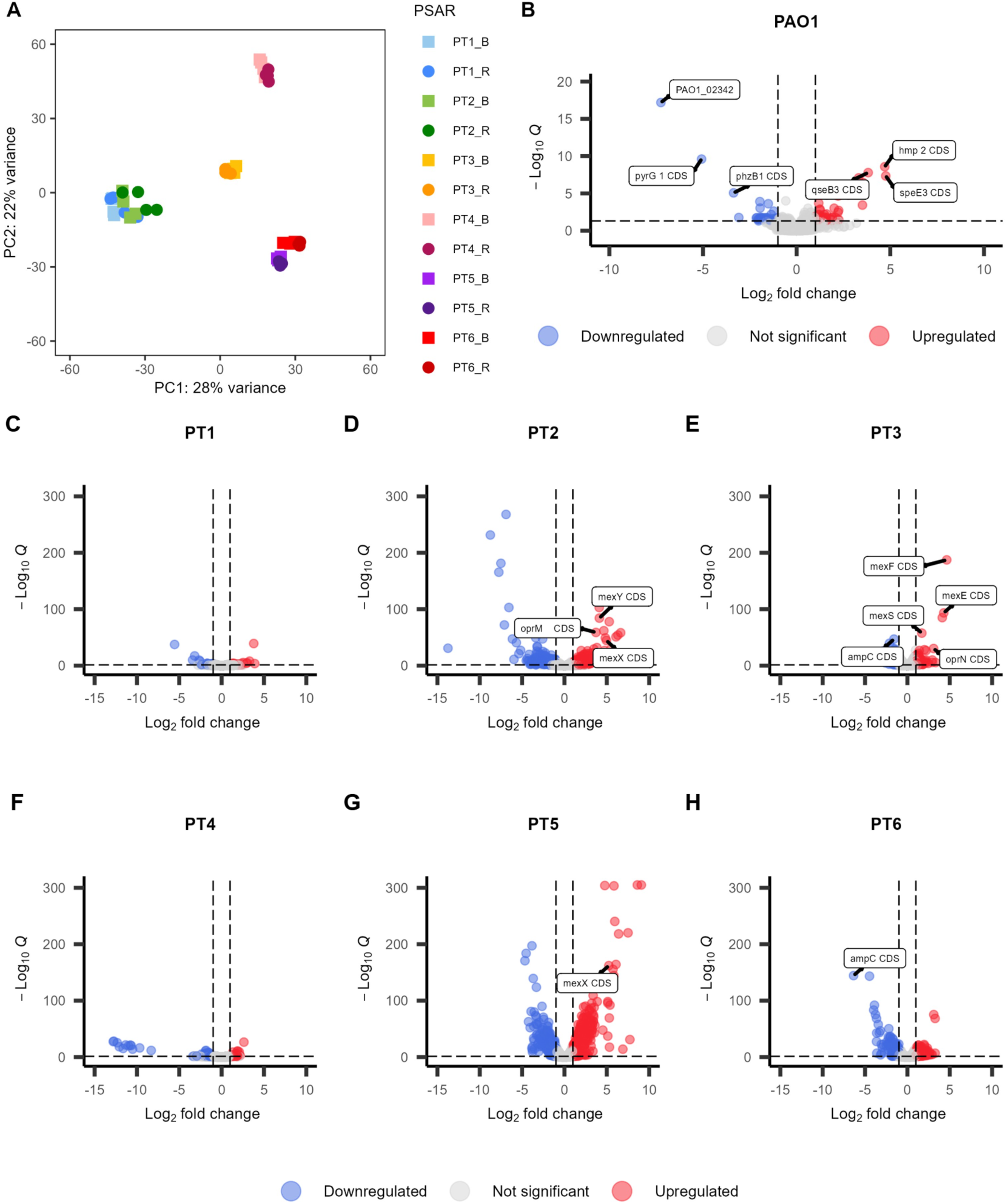
Gene expression differences between resistant and baseline isolates of all six patients against PAO1 and each other. (A) PCA plot of PAO1 against all baseline and resistant patient isolates. Each dot represents one biological replicate. (B) Volcano plot depicting differentially regulated genes of all baseline vs all resistant isolates. (C-H) Volcano plot of baseline vs resistant isolate of each individual patient where only significant DEGs associated with Imi/Rel resistance and other efflux-pump related genes are labelled. The x-axis shows log2 fold change in expression level, and the y-axis shows the significance of the difference in expression. Analysis in panels A and B used PAO1 as background to be compared to. For DESeq2 analysis in panel B, design = ∼0 + condition.

To further explore transcriptomic changes within each patient, we used the baseline isolate from each patient as a reference for read mapping and performed PCA analysis on the transcriptomes of the baseline and resistant isolates from each patient (**Fig S2**). The resulting plots showed that the expression profiles of each patient’s resistant isolates were distinct from that of the paired susceptible isolates. Notably, the number, magnitude, and identity of DEGs varied considerably between patients (**Fig 1C-H**). Isolates from PT1 showed the fewest and smallest changes (n=80 DEGs) while isolates from PT2 had the most extensive transcriptomic response (n=949 DEGs) (**File S1**). Among the DEGs from four patients (PT2, PT3, PT5, and PT6), we identified several genes belonging to previously characterized Imi/Rel resistance pathways (**Fig 1D-E, G-H, File S1**) (5, 7). Specifically, Imi/Rel-resistant isolates from PT2, PT3, and PT5 showed upregulation of efflux pump-related genes (*mexXY-oprM, mexEF-oprN, mexS*) relative to the corresponding baseline isolates (**Fig 1D-E, G**). For these isolates, efflux pump overexpression was likely induced by Imi/Rel exposure and might result in reduced intracellular concentrations of the antibiotic (22). Post-exposure isolates from PT3 and PT6 also showed downregulation of *ampC* expression (**Fig 1E, H**). This contrasts with previous reports of *ampC* overexpression following imipenem exposure, suggesting that the combination with relebactam may differentially influence *ampC* expression (23). Interestingly, PT1 and PT4 showed no DEGs related to efflux pumps or *ampC* (**Fig 1C, F**), suggesting that there may be additional, uncharacterized transcriptional mechanisms involved in *P. aeruginosa* Imi/Rel resistance.

To assess whether DEGs were enriched for particular functions, we assigned each gene in the baseline isolate genome as well as each DEG to a Clusters of Orthologous Genes (COG) category. Among the DEGs identified as upregulated or downregulated between baseline and Imi/Rel-resistant isolates from PT2 and PT5, we observed significant enrichment of upregulated genes related to COG Category J (Translation, Ribosomal Structure, and Biogenesis), while the same comparison between isolates from PT6 showed significant downregulation of genes in this category (**Fig 2; File S1**). The Imi/Rel-resistant isolate from PT3 showed significant upregulation of genes related to Category Q (Secondary Metabolite Biosynthesis, Transport and Catabolism), while downregulated genes in the Imi/Rel-resistant isolate from PT5 were enriched in Category E (Amino Acid Transport and Metabolism) and Category S (genes of Unknown Function) (**Fig 2; File S1**). Finally, the Imi/Rel-resistant isolate from PT6 had significant downregulation of genes in Category O (Post-translational Modifications) (**Fig 2; File S1**). Collectively, these results indicate that treatment-emergent Imi/Rel resistance in *P. aeruginosa* could be associated with broader transcriptional changes affecting other metabolic processes besides antibiotic efflux and *ampC* expression.

**Figure 2.**
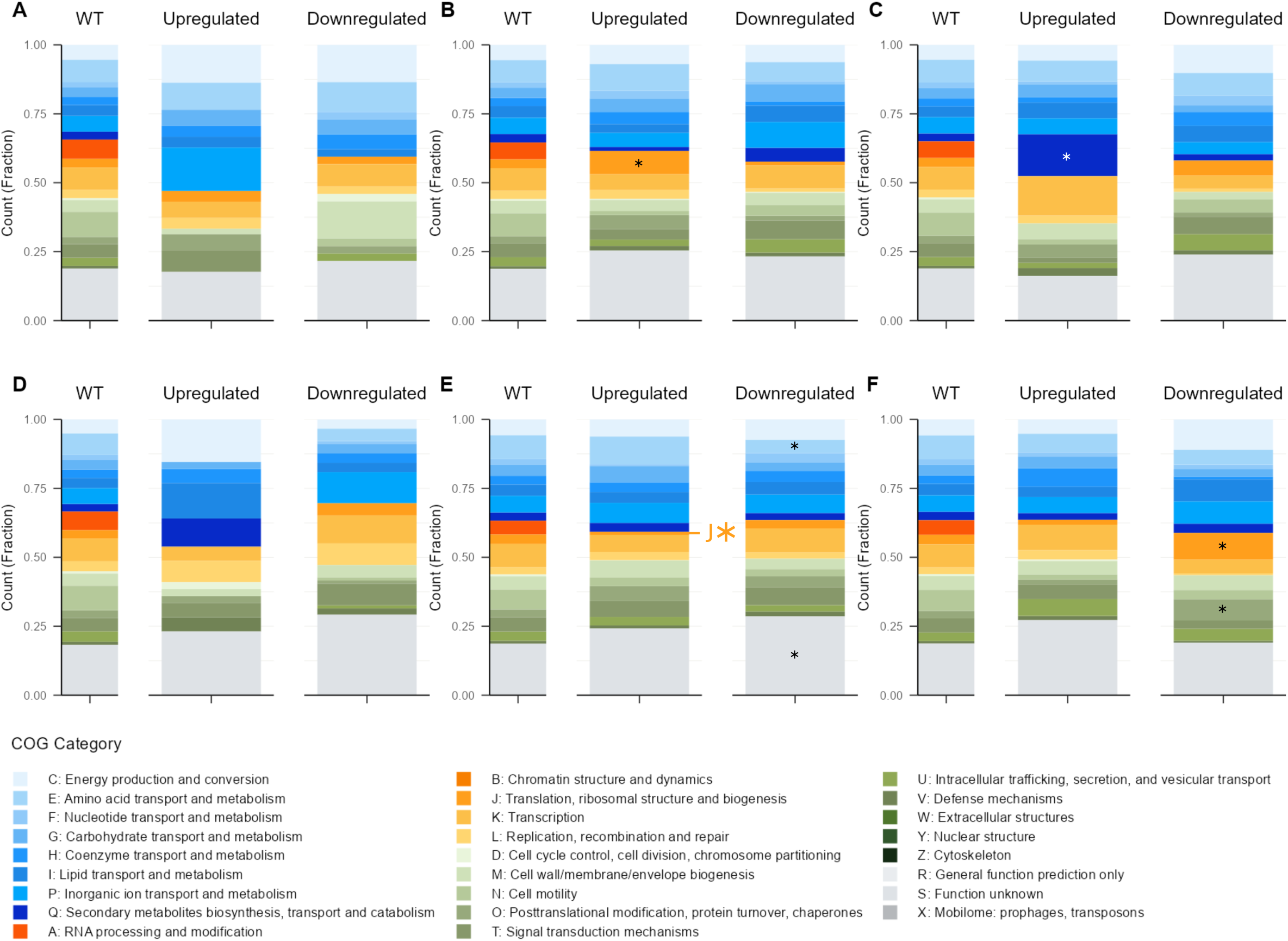
Distribution of COG categories in isolate genomes and among differentially expressed genes in post-exposure isolates for each patient. PT1 (A), PT2 (B), PT3 (C), PT4 (D), PT5 (E), PT6 (F). COG categories were assigned using EggNOG-mapper and genes that did not have assigned COG categories were excluded. The distribution of differentially expressed genes in each resistant isolate was compared with the distribution of all genes in each baseline isolate genome using Fisher’s exact test with Bonferroni correction. *pS0.0027.

The top 10 up- and downregulated genes between pre- and post-exposure isolates, based on expression fold change, were then compared to each other, with equivalent genes present across isolates determined through NCBI BLAST using a >90% identity cut-off (**Fig 3A, B**). Top DEGs were unique to each patient, with one notable exception; the phenazine biosynthesis gene *phzB1* was downregulated in post-exposure isolates from both PT2 and PT3. We also observed several genes that were upregulated in the post-exposure isolate from one patient but downregulated in another patient, including a hypothetical protein in PT2 and PT6, the transcriptional regulator *dmlR_12* and putative cysteine hydrolase *ycaC_3* in PT3 and PT1, and the carbazole-degradation related hydrolase *carC* in PT4 and PT6 (**Fig 3A, B**) (24–26). Taken together, these findings further highlight the diverse and background-specific effects of treatment-emergent Imi/Rel resistance in *P. aeruginosa*.

**Figure 3.**
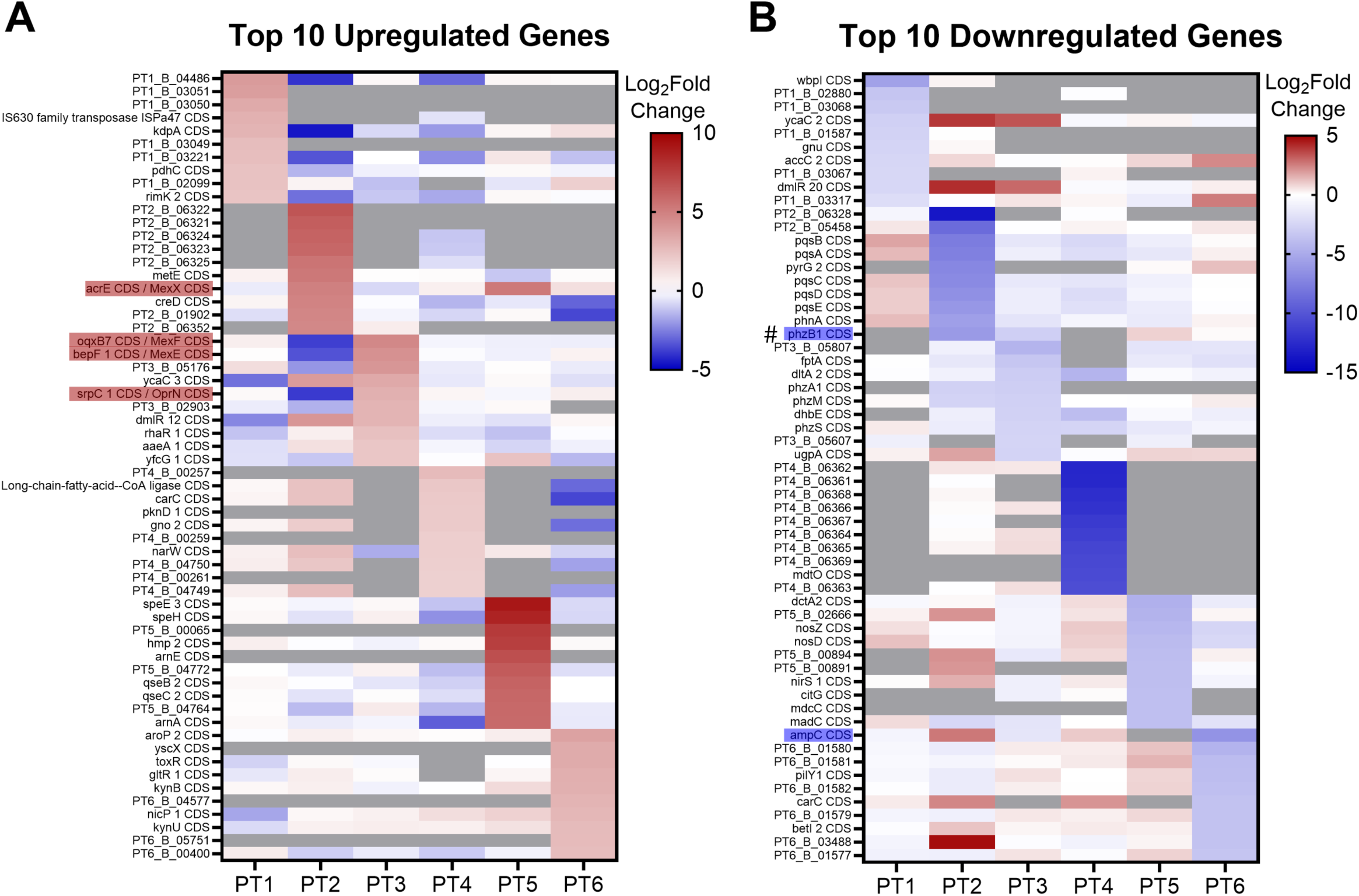
Top 10 upregulated and downregulated genes in all six patients. The top 10 genes that were differentially regulated are unique in all patients, except for one gene that are shared by PT2 and PT3 (denoted by a pound sign). *phzB1, ampC,* and *mex* operon genes are shaded in either blue or red. Grey color denotes genes that are not present in that patient or with no expression data.

## Discussion

In this study we investigated transcriptional changes among *P. aeruginosa* clinical isolates associated with treatment-emergent resistance to the β-lactam/β-lactamase inhibitor combination Imi/Rel. Comparing all pre-exposure isolates to all post-exposure isolates initially suggested a conserved transcriptomic response accompanying Imi/Rel resistance. However, this signal was not seen among individual patient transcriptional profiles, suggesting significant changes in a subset of isolates rather than a common response across all *P. aeruginosa* isolates. Interestingly, these shared responses did not include previously characterized Imi/Rel resistance mechanisms, except for *phzB1,* a phenazine biosynthesis protein involved in *P. aeruginosa* biofilm formation and antibiotic tolerance, which was downregulated in isolates from two patients (27, 28). Individual per-patient analysis revealed that both individual DEGs and functional categories of DEGs were highly variable across different patients. Isolates from only four of the six patients showed transcriptional changes in known Imi/Rel resistance-associated genes, suggesting that *P. aeruginosa* may achieve Imi/Rel resistance through other unknown pathways.

Overall, the fact that we did not identify strongly conserved transcriptional changes among post­exposure isolates indicates that evolutionary pathways to Imi/Rel resistance likely vary across genetically distinct isolates. Future studies that include additional patients, and/or examine how DEGs affect Imi/Rel susceptibility through functional validation studies, are warranted. Such efforts would inform the optimized use of Imi/Rel to treat MDR *P. aeruginosa* infections.

## Author Notes

Two supplemental figures and two supplemental files are available with the online version of this article.

## Abbreviations

COG: Clusters of Orthologous Genes
COG: DEG, differentially expressed gene
Imi/Rel: Imipenem-relebactam
LVAD: Left-Ventricular Assist Device
MDR: multidrug-resistant
MIC: minimum inhibitory concentration
PT: patient
ST: Sequence type

## Acknowledgements

This work was funded by an investigator-initiated grant from Merck & Co. awarded to RKS and DVT.

## Data Availability

Raw RNA sequencing reads generated in this study are available under GEO project GSE343731. Whole genome sequencing data are available under BioProject PRJNA1512454.

## Ethics Approval

The study was approved by the University of Pittsburgh Institutional Review Board (STUDY22070065) with a waiver of informed consent.

## Supplemental Material

**Figure S1.**
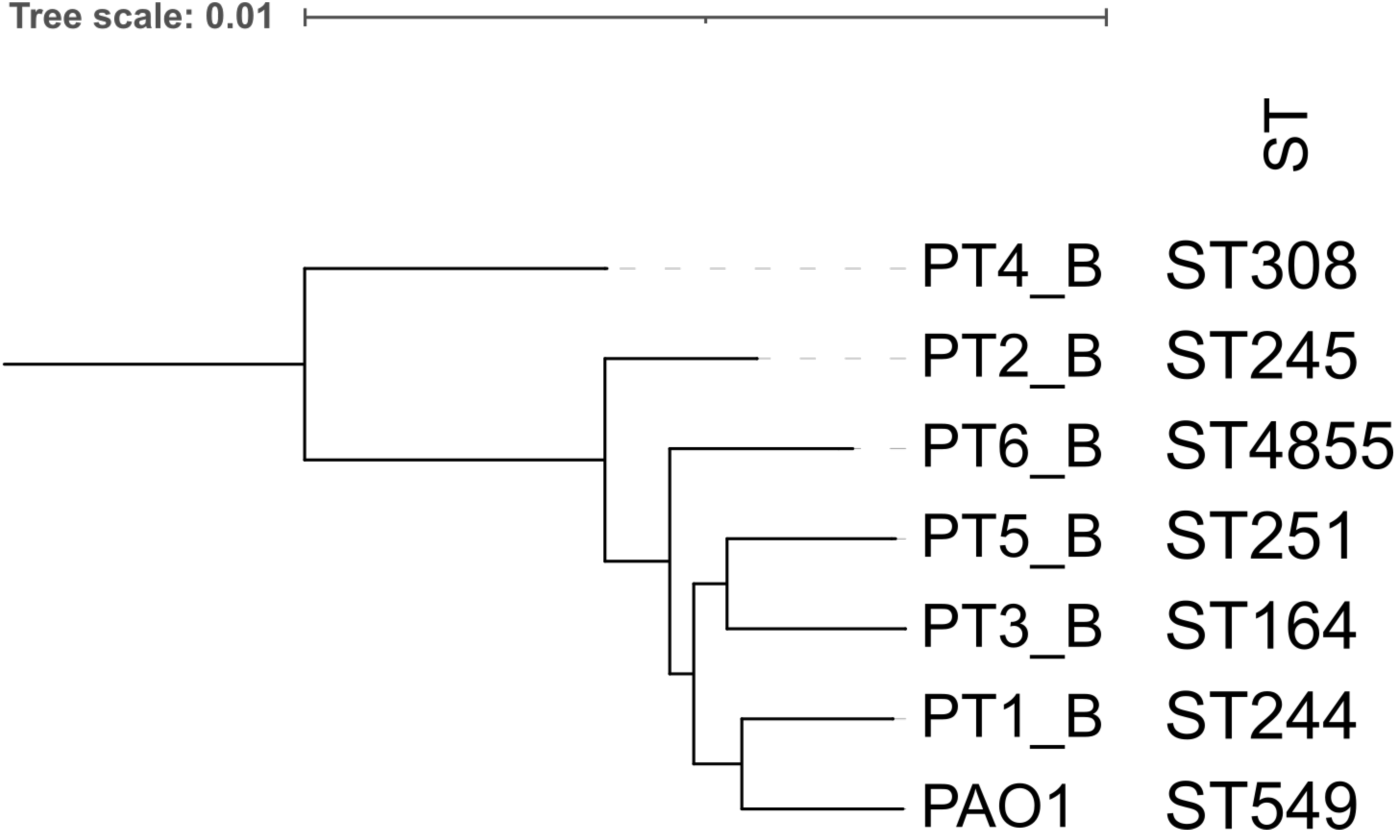
Core-genome phylogeny tree of pre-exposure isolates and PAO1. Core genome phylogenetic tree was created using RaxML and visualized with iToL. Sequence types (STs) were determined by PubMLST.

**Figure S2.**
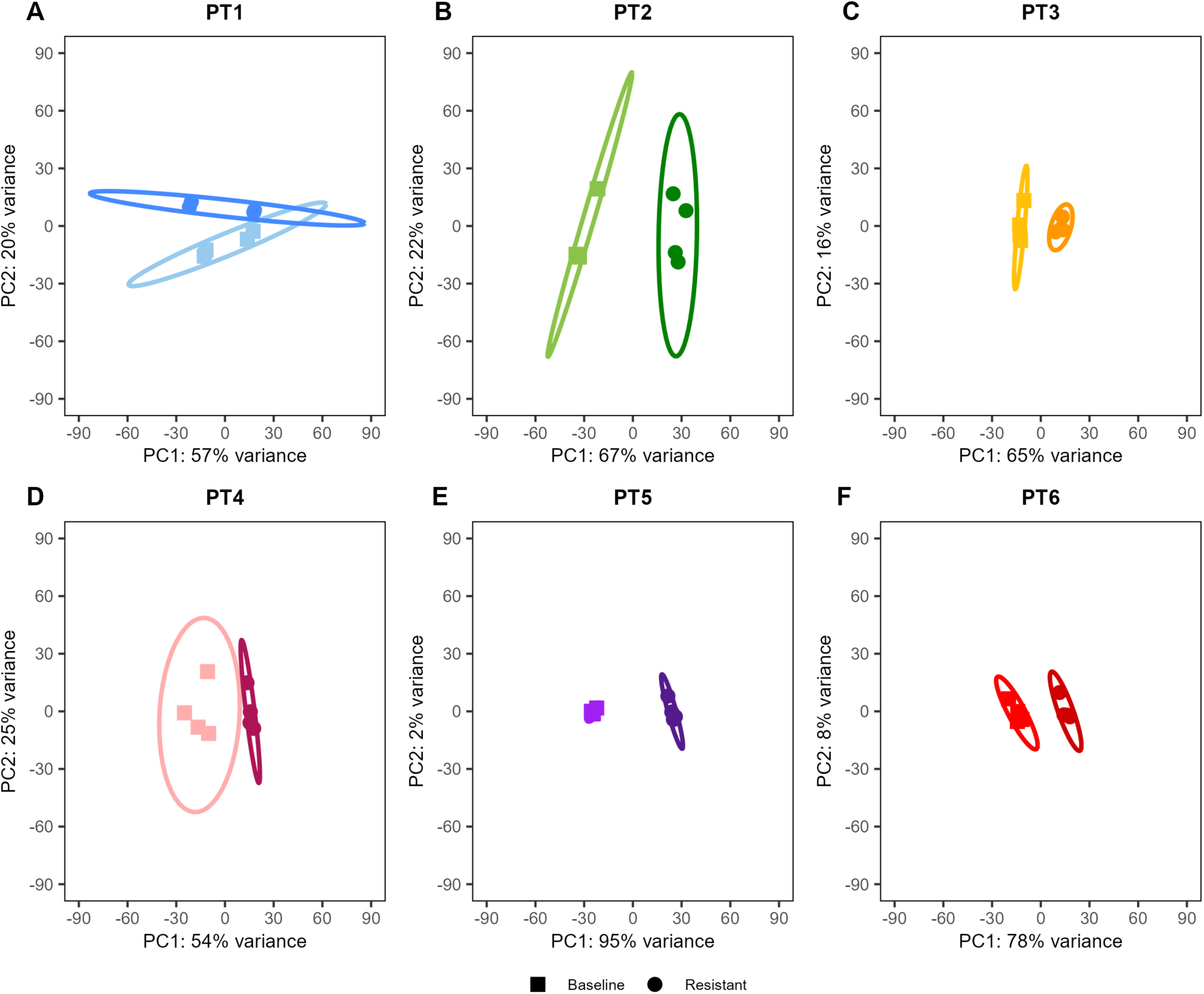
Individual PCA plots based on transcriptional profiles of pre- and post-exposure isolates from each patient. Expression data from each patient’s resistant isolate was compared to the data from the patient’s baseline isolate. Expression data of the biological replicates from all patient conditions cluster according to their condition (baseline vs resistant). Ellipses represent 95% confidence interval. Each dot represents one biological replicate.

**File S1.** Raw means of DESeq2 Analysis and COG category assignment.

**File S2.** Top 10 differentially regulated genes from all six patients.

